# Reducing Supply Chain Dependencies for Viral Genomic Surveillance: Get by with a Little HELP from Commercial Enzymes already in your Lab Freezer

**DOI:** 10.1101/2025.06.11.658579

**Authors:** Ganna Kovalenko, Myra Hosmillo, Chris Kent, Kess Rowe, Andrew Rambaut, Nicholas J Loman, Joshua Quick, Ian Goodfellow

## Abstract

The COVID-19 pandemic exposed vulnerabilities in global laboratory supply chains, disrupting genomic surveillance efforts essential to epidemic response. To address this challenge, we developed ARTIC HELP (Homebrew Enzymes for Library Preparation), a practical, open-source adaptation of the widely adopted ARTIC nanopore sequencing protocol for viral genomic surveillance. We describe generic, cost-effective alternatives to all enzyme mixes used in tiling multiplex RT-PCR amplification of the virus genome, and the nanopore native barcoding workflow, including end-prep (EP), barcode ligation (BL), and adapter ligation (AL), making it broadly applicable to any laboratory. Through systematic evaluation, we identified a wild-type M-MLV reverse transcriptase and two types of proofreading DNA polymerases as effective alternatives when standard reagents are unavailable due to high cost or limited supply: B-family Pfu-based polymerases with a fused Sso7d DNA-binding domain, and blends combining A-family (Taq-based) and B-family (Pfu-based) polymerases. Validation on clinical samples of SARS-CoV-2 and Norovirus GII confirmed that the HELP workflow achieves genome coverage comparable to the ARTIC LoCost protocol. For SARS-CoV-2 samples (Ct ≤28), the wild-type M-MLV RT combined with selected Pfu or A+B polymerases, along with optimised HELP mixes (EP, BL, AL), achieved genome coverage of 84.0–99.6%. For Norovirus GII (Ct ≤32), the HELP workflow using one of the Pfu polymerases achieved genome coverage of >85% for six out of eight genotypes tested. Notably, several of the other polymerases tested showed reduced performance at higher Ct values. However, they still achieved strong coverage at Ct <24, supporting their use as emergency alternatives in rapid outbreak-response sequencing when viral input is high and RNA quality is sufficient. Our approach, ARTIC HELP, provides a framework which can be implemented to address supply chain disruptions, while maintaining robust genomic sequencing capabilities. A cost analysis highlights the well-known significant global disparities in reagent pricing, driven not by protocol differences but by import fees and supply barriers. Thus, our findings highlight the need for fairer global pricing models and support for local sourcing strategies like HELP, to promote equity in genomic research and ensure preparedness for future public health challenges.

## Introduction

When the West African Ebola outbreak hit in 2014, sequencing a virus in real time, let alone in on-site settings, was effectively unprecedented in outbreak settings (*1*). That crisis marked a turning point, driving the development of genomics-informed approaches to global pathogen surveillance systems that have since reshaped public health responses to emerging infectious disease threats (*2, 3*). The COVID-19 pandemic reinforced this shift, catalysing a wave of investment in global sequencing infrastructure and prompting the World Health Organization to launch a 10-year strategy (2022–2032) to expand genomic surveillance capacity worldwide (*4-6*). These efforts laid the foundation for integrating genomics into routine public health practice, supporting timely detection and response to infectious disease outbreaks.

One of the most influential efforts that emerged in response to this shift was the ARTIC Network, the first coordinated initiative to deliver end-to-end, on-site deployable protocols for portable genomic surveillance (https://artic.network/ ; https://community.artic.network/t/a-beginners-guide-to-artic/531). Launched during the Ebola outbreak and aligned with the rise of Oxford Nanopore Technology (ONT) sequencing, ARTIC has since developed tools for rapid viral genome sequencing and analysis, enabling real-time response in real-world outbreak settings. The network has played a pivotal role in responses to Zika, SARS-CoV-2, and more recently, Monkeypox (*7-9*). Drawing on experience from Ebola and Zika, the ARTIC team rapidly designed and released a whole-genome sequencing protocol for SARS-CoV-2 at the onset of the pandemic. The ARTIC primers, based on an amplicon tiling scheme, became the backbone of SARS-CoV-2 genomic surveillance, contributing to over 18 million genome sequences across platforms and enabling timely variant detection and public health response (*10-13*). To support broader adoption and higher-throughput use, the ARTIC LoCost workflow was introduced, reducing reagent volumes and lowering costs (*14, 15*). This cost-effective protocol has been widely implemented in laboratories worldwide. More recently, adaptations of the ARTIC workflow have also been applied to wastewater surveillance and other viral targets, demonstrating its flexibility and ongoing relevance in public health genomics (*13, 16*).

Yet the pandemic also exposed the limitations of even widely adopted protocols, revealing structural weaknesses in global diagnostic and genomic surveillance systems, particularly their reliance on protocols that depend on specific reagents without validated alternatives. This lack of flexibility made laboratory operations highly vulnerable to supply chain disruptions, leading to shortages and delays in essential reagents, enhanced by transportation restrictions and increased competition for resources (*17, 18*), making even widely used protocols such as ARTIC difficult to sustain. To remain effective in future outbreaks, genomic workflows must be not only technically robust but also adaptable to resource constraints and resilient in the face of supply chain instability. While several high-level strategies have been proposed to improve resilience in health-related supply chains, such as supply diversification, local sourcing, and improved logistics, these efforts primarily focus on procurement and distribution systems, rather than the design of laboratory workflows themselves (*17*). Our work addresses this under-served layer by filling a precise, practical gap: the need for validated, protocol-level flexibility in genomic surveillance workflows.

In response to these challenges, we developed and validated the ARTIC HELP workflow (Homebrew Enzymes for Library Preparation) as a flexible alternative to the ARTIC LoCost protocol. We designed HELP to include open-source substitutes for all key enzymatic steps in the native barcoding workflow (ONT), including end-repair, barcode ligation, and adapter ligation, making the workflow broadly applicable to any laboratory using nanopore sequencing. Rather than aiming to replace commercial enzyme mixes entirely, we focused on providing practical alternatives that use enzymes and buffers commonly found in standard molecular biology laboratories. We validated the HELP workflow on clinical samples of SARS-CoV-2 and Norovirus GII, confirming its performance under real-world conditions. By offering reliable substitutes for critical reagents, we aim to strengthen the resilience and continuity of pathogen genomic surveillance during public health crises.

## Materials and methods

### Virus culture

Live SARS-CoV-2 (SARS-CoV-2/human/Liverpool/REMRQ001/2020) was cultured in Vero-E6 (ATCC) cells grown in Dulbecco’s Modified Eagle Medium (Pan Biotech) supplemented with 1% Glutamine (Gibco), 10% foetal calf serum, 100 U/ml penicillin and 100 μg/ml streptomycin (Thermo Scientific), at 37 °C with 5% CO2.

### Viral RNA extraction

SARS-CoV-2 control panel RNA was generated using the isolate SARS-CoV-2/human/Liverpool/REMRQ001/2020. Following SARS-CoV-2 live virus inoculation, infected cells were collected by centrifugation at 12000x g for 5 mins. Cells were lysed in lysis buffer containing 4M guanidine isothiocyanate, 1% beta-mercaptoethanol and 2% Triton X-100 for 10 mins at room temperature. After lysis, samples were centrifuged through a QIAshredder homogenizer and ethanol was then added to the cleared lysates. Total RNAs were isolated and purified using the Sigma GenElute protocol, and the viral RNA was eluted with 60 μl of RNase-free water.

Stool specimens were originally obtained and anonymized with written consent from patients at Addenbrooke’s Hospital in Cambridge, United Kingdom, who tested positive for HuNoV infection. These samples were collected under the ethical approval (REC-12/EE/0482) for a previous study (*19*). Each specimen was diluted 1:5 or 1:10 (wt/vol) with phosphate-buffered saline (PBS) depending on the water content of the stool sample. Diluted samples were then vortexed vigorously for 30 sec and centrifuged for 5 min at 8000 x g at room temperature. Following centrifugation, 200 μl aliquots of the supernatants were collected for total RNA extraction or immediately frozen at -80°C until required. RNA was extracted from the samples using the Sigma GenElute protocol using 700 μl of guanidine isothiocyanate (GITC)-containing buffer with 1% beta-mercaptoethanol per 200 μl of diluted stool samples. The RNA was purified according to the manufacturer’s instructions and the viral RNAs were eluted with 30 μl of RNase-free water.

### RT-qPCR for the detection and quantification

Viral RNA levels in all samples were confirmed immediately prior to use. Ct-values were determined by RT-qPCR for SARS-CoV-2 using a standard diagnostic workflow for 2019-nCoV screening (*20*) and for Norovirus GII (*21*). A ten-fold serial dilution of *in vitro* transcribed RNA was used to generate a standard curve and determine the absolute copy number of viral RNA.

### cDNA synthesis and multiplex PCR

Detailed master mix recipes and a full list of reagents including catalog numbers used in the HELP study are provided in the Extended Data (Table S1 and Table S2). The ARTIC LoCost sequencing protocol for SARS-CoV-2 v3 was used as the baseline (*15*).

M-MLV reverse transcriptase (Promega) was explored as an alternative to the standard LunaScript RT (NEB) used in the ARTIC LoCost sequencing protocol. For cDNA synthesis, a reaction comprising of random hexamers (Invitrogen), dNTP Mix (Thermo Fisher), RNase OUT (Invitrogen), and 12 μL of template RNA was prepared (*Extended data:* Table S1). The RT reaction was carried out under the following conditions: 25°C for 5 min, 42°C for 50 min and 70°C for 10 min. Six DNA polymerases were used alongside the standard Q5 Hot Start High-Fidelity DNA Polymerase (NEB), their characteristics are summarised in Extended data Table S3. These included: 1) (Platinum) Platinum SuperFi DNA Polymerase (Invitrogen); 2) (PrimeSTAR) PrimeSTAR GXL DNA Polymerase (TAKARA); 3) (KAPA) KAPA Taq Extra HotStart ReadyMix PCR Kit (KAPAbiosystems); 4) (EcoDry) High Fidelity PCR EcoDry Premix (TAKARA); 5) (Phusion) Phusion High-Fidelity DNA Polymerase (Thermo Scientific); 6) (KOD) KOD Hot Start DNA Polymerase (Sigma-Aldrich). Multiplex amplicon-based PCR was run using artic-sars-cov2/400/v4.1.0 (*22*) and norovirus-gii/800/v1.1.0 (*23*) primer schemes. Then 2.5 μL of cDNA was added to the PCR reactions for each pool, bringing the total reaction volume to 25 μL (*Extended data:* Table S1). Samples were amplified under the same PCR cycling conditions, adapted from the LoCost protocol (*15*): heat activation at 98°C for 30 seconds, followed by 30 cycles of 95°C for 15 seconds and 63°C for 5 minutes. The PCR reactions were then pooled together, purified, and quantified following the LoCost protocol before end-prep. Briefly, 5 μL of each PCR reaction from each pool were combined in 40 μL of nuclease-free water (NFW) and purified using a 1:1 ratio of PCRClean DX magnetic beads (Aline BioSciences), then quantified using the Qubit dsDNA High Sensitivity assay (Life Technologies) on the Qubit Flex fluorometer. In some instances, gel electrophoresis (1% agarose) was performed to analyse the PCR products generated by each condition.

### Library preparation (end-prep, barcode and adapter ligation)

#### End-prep (EP)

The ends prep reaction incorporates a number of enzymatic reactions; the ends of the amplicons are first polished using the Klenow fragment of DNA Polymerase I (Klenow) (NEB) and T4 DNA Polymerase (NEB), then phosphorylated and adenylated using T4 Polynucleotide Kinase (T4 PNK) (NEB) and Taq DNA Polymerase (Thermo Scientific), respectively. Three options of HELP EP mixes were tested (A, B and C), which varied in the concentrations of enzymes described in more detail in the text (*Extended data:* Table S1). All mixes included 1X T4 DNA Ligase Reaction Buffer (50 mM Tris-HCl, 10 mM MgCl2, 1 mM ATP, 10 mM DTT), 0.5 mM dNTPs, and 5% PEG-8000 for efficiency. Reactions contained 3.3 μL of amplicons, with NFW to a final volume of 10 μL. The reaction was incubated for 30 minutes at 20°C, 30 minutes at 65°C, and finally cooled on ice.

#### Barcode Ligation (BL)

The HELP BL master mixes were developed as alternatives to the NEBNext® Ultra™ II Ligation Module and Blunt/TA Ligase Master Mix (NEB), and are detailed in Extended Data Table S1. Five formulations (A–E) were prepared using T4 DNA Ligase at 1000 U per reaction, along with ligation enhancers including hexamine cobalt chloride (HCC), 1,2-Propanediol (1,2-PrD), PEG-8000, and two concentration options of the T4 DNA Ligase Reaction Buffer (NEB). The native barcoding expansion kit (EXP-NBD196, ONT) was used, with barcodes diluted in a ratio of 1.4:1 with NFW, prior to use (NFW:barcode). Subsequently, 3 μL of the diluted barcodes were utilized per reaction. For each reaction, 1.5 μL of end-prepared amplicons from the EP step were subjected to barcode ligation. NFW was added to adjust the reaction volume to 10 μL, and the reaction was incubated for 30 minutes at 20°C, 10 minutes at 70°C, and then cooled on ice.

#### Adapter Ligation (AL)

Following barcoding, individual reactions were pooled, purified using a 0.4× volume of PCRClean DX magnetic beads, and quantified by Qubit, in line with the ARTIC LoCost protocol. Adapter ligation was then performed using alternative HELP AL mixes developed in place of the NEBNext® Quick Ligation Module (NEB) (*Extended data:* Table S1). The AL reaction was set up in 2X T4 DNA Ligase Reaction Buffer (100 mM Tris-HCl, 20 mM MgCl_2_, 2 mM ATP, 20 mM DTT, pH 7.5) and 10 % of PEG-8000. Two HELP AL formulations were tested: HELP AL-A (4000 U T4 Ligase/reaction) and HELP AL-B (2000 U T4 Ligase/reaction). Each reaction included 30 μl of the barcoded amplicon pool and 5 μl of Adapter Mix II (AMII, EXP-NBD196, ONT), with the final volume adjusted to 70 μl using NFW. For comparison, a parallel adapter ligation using the commercial Rapid DNA Ligation Kit (Thermo Scientific) was set up in a 50 μl reaction volume. All ligation reactions were incubated at room temperature for 20 minutes. Libraries were then cleaned using a 1:1 ratio of PCRClean DX beads and quantified according to the LoCost protocol.

### Sequencing and Bioinformatics workflow

Final libraries were generated using the Ligation Sequencing Kit (SQK-LSK109, ONT), loaded onto FLO-MIN106 flow cells on a GridION device (ONT) and sequenced using the MinKNOW software using real-time basecalling with a high accuracy model. Demultiplexing was conducted using the barcode-both-ends option and read filtering based on quality score 9. The ARTIC nCoV-2019 novel coronavirus bioinformatics protocol using reference-based medaka workflow was used to process the output into consensus genome sequences (*24*). Genome regions with depth of <20× coverage were masked and represented with N characters. The sequencing results were additionally analysed using ncov-tools (*25*). For Norovirus GII sequences the ARTIC field bioinformatics pipeline was applied using reference-based medaka workflow (*26*). The closest reference genomes were selected using Rampart (*27*). Norovirus sequences were analysed with the RIVM Norovirus Typing Tool v.2.0 (*28*). Data plots and heatmaps were generated using Jupyter Notebook (https://jupyter.org/).

## Results

The study was conducted in three stages (Figure 1). In Stage 1, we systematically replaced reagents at each step of the ARTIC LoCost sequencing protocol across six workflows (wf), each targeting one of the five experimental steps: RT, PCR, and the library preparation steps (EP, BL, and AL). This approach aimed to identify viable reagent alternatives. In Stage 2, we determined the optimal replacements within a single library workflow, ensuring the new components worked seamlessly together. Sequencing performance in Stages 1 and 2 was tested across four SARS-CoV-2 RNA concentrations from the lab-grown isolate (SARS-CoV-2/human/Liverpool/REMRQ001/2020), corresponding to Ct values 21.2, 24.6, 27.9, and 31.4, equating to 2.2×10^4, 2.01×10^3, 1.83×10^2, and 1.65×10^1 RNA copies/reaction, respectively. Finally, Stage 3 validated the best replacements using clinical samples of SARS-CoV-2 and Norovirus GII (with Ct-values <33), confirming their effectiveness in real-world applications. HELP workflows from Stage 3 are available on protocol.io https://protocols.io/view/artic-help-protocol-for-amplicon-based-viral-genom-gzsibx6cf. Performance metrics included read count, percentage of mapped reads, mean and medium depth, genome coverage, and amplicon dropouts (regions with ≤20x depth), detailed in Extended data Tables S4–S6.

**Figure 1.**
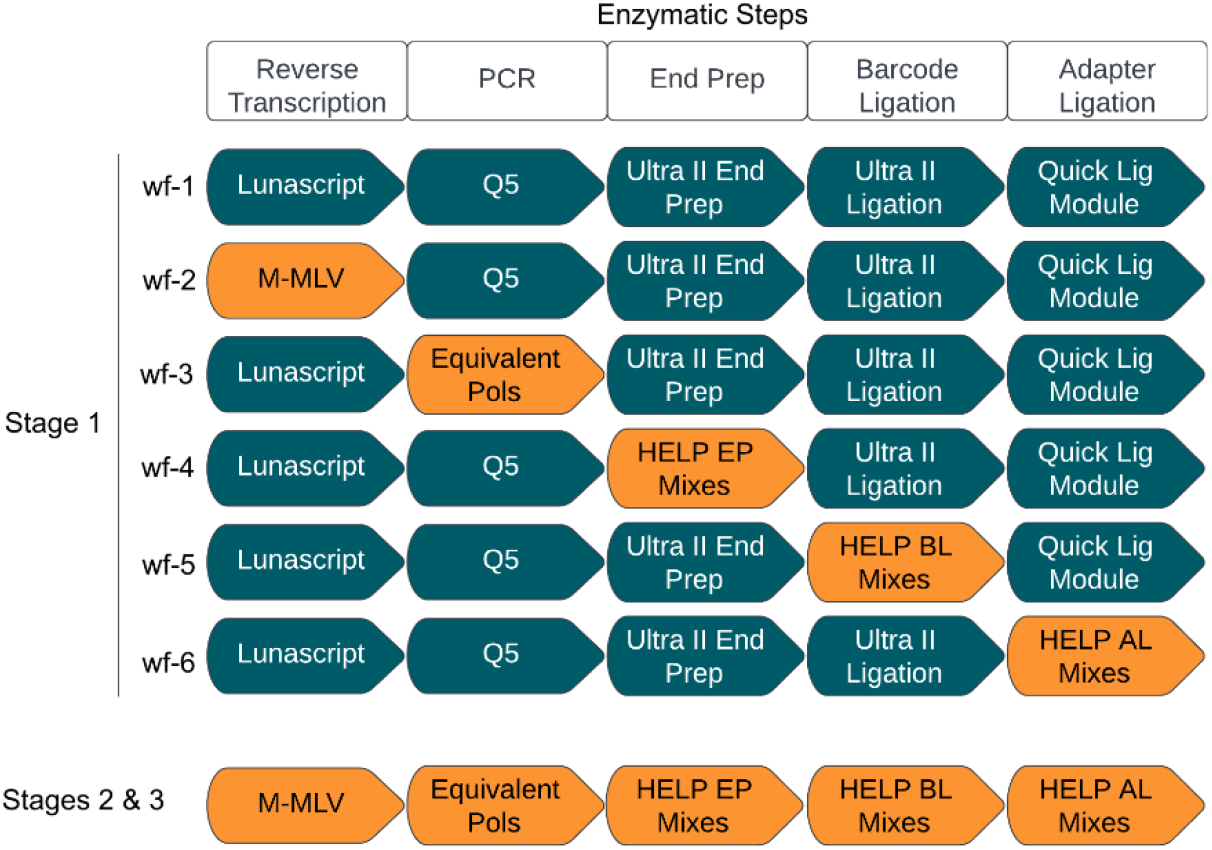
Schematic representation of the HELP study stages. Stage 1 highlights the six workflows (wf) designed to assess the efficacy of generic enzyme replacements across the various enzymatic steps in the ARTIC LoCost workflow. Workflow 1 (wf-1) represents the reference ARTIC LoCost protocol where all commercial mixes are used. In workflows 2 to 6 (wf-2 to wf-6), reagents were systematically replaced at each enzymatic step in the library preparation process using generic equivalents (highlighted in orange and described in more detail in the text). Stages 2 and 3 represent the evaluation of the most effective replacements identified in Stage 1 within a single library preparation workflow.

### Stage 1: Systematic replacement of the core reagents in each step of the ARTIC sequencing workflow

Workflow 1 (wf-1) followed the ARTIC LoCost protocol using commercial mixes from NEB, while workflows wf-2 to wf-6 systematically tested alternative enzyme mixes at each step of library preparation. Figure 2 summarizes this comparison, showing genome coverage across four Ct values using a SARS-CoV-2 isolate. In wf-2, LunaScript RT was replaced with M-MLV RT, while the remaining steps utilised reagents from the LoCost protocol. In wf-3, Q5 DNA Polymerase was replaced with one of six alternatives: Platinum, PrimeSTAR, KAPA, EcoDry, Phusion, or KOD. By gel electrophoresis, we assessed the amplification efficiency from wf-1, wf-2 and wf-3 (*Extended data:* Figure S1) and found that while all polymerases generated expected products, amplification efficiency varied. KOD polymerase was particularly inefficient and excluded from further experiments. We also noted that EcoDry, PrimeSTAR, and KAPA produced longer (∼1,000 bp) chimeric products in samples with lower Ct values, a known artifact in PCR amplifications (*29*).

**Figure 2.**
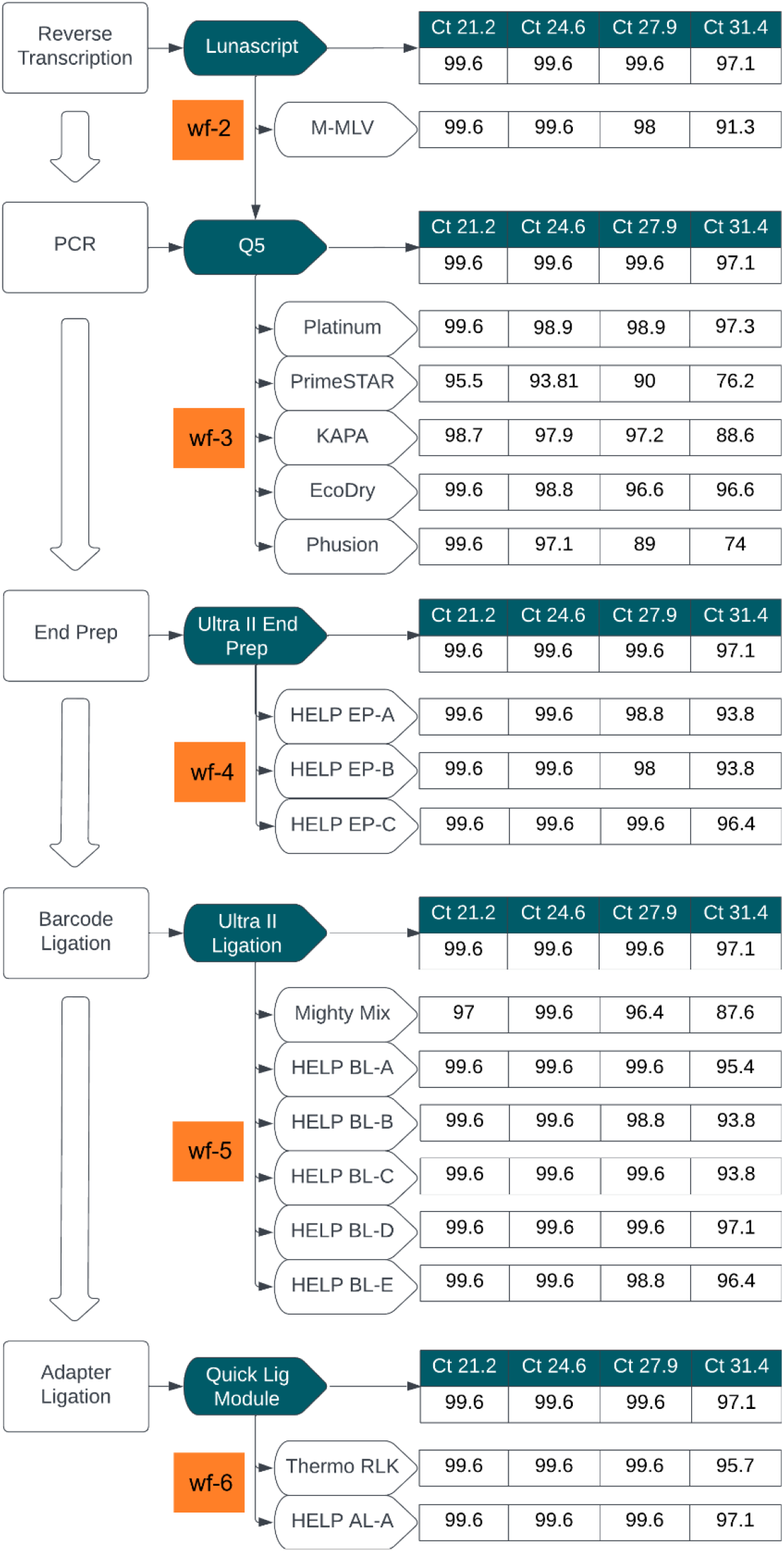
Stage 1: Summary of workflow comparisons. Flowchart summarising the workflow comparison for library preparation, evaluating alternative enzyme mixes at key steps in the LoCost protocol. Genome coverage percentages obtained using SARS-CoV-2 isolate across four Ct values (21.2, 24.6, 27.9, and 31.4) is shown for each workflow. The reference ARTIC LoCost protocol (wf-1) is represented by the reagents in dark green boxes, and the % genome coverage obtained is listed first for each step as the reference. Alternative workflows (wf-2 to wf-6) are highlighted in orange, corresponding to specific enzymatic steps tested in reverse transcription, PCR, end prep, barcode ligation, and adapter ligation.

Replacing LunaScript with M-MLV RT combined with Q5 polymerase (wf-2) delivered comparable performance to the ARTIC LoCost protocol, except at the highest Ct value (31.4), where M-MLV showed reduced genome coverage (91.3%) and 10 amplicon dropouts (Figures 2 and 3). In contrast, the LoCost workflow (LunaScript) maintained 97.1% coverage with only three dropouts, highlighting the importance of RT enzyme selection. Nonetheless, M-MLV RT demonstrated viability as an effective alternative, achieving comparable genome coverage and sequencing depth under most conditions.

**Figure 3.**
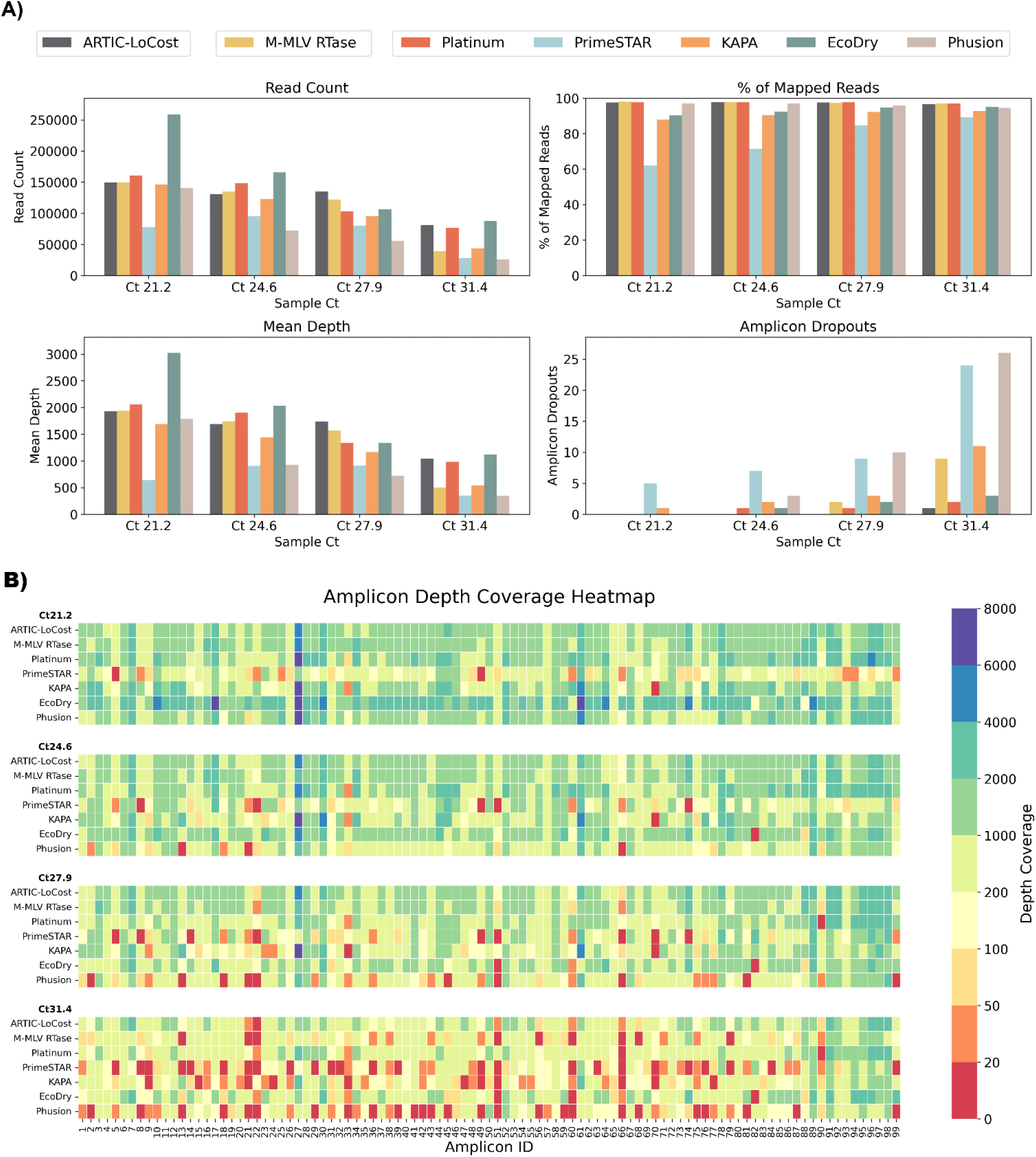
Stage 1: Comparison of M-MLV RTase and alternative polymerases to the LoCost protocol. **(A)** Bar plots comparing the performance of M-MLV RTase (wf-2) and five polymerases,Platinum, PrimeSTAR, KAPA, EcoDry, and Phusion (wf-3),against the LoCost protocol (wf-1) across four key metrics: Read Count, % of Mapped Reads, Mean Depth, and Amplicon Dropouts. Evaluations were conducted using SARS-CoV-2 isolate RNA at Ct values of 21.2, 24.6, 27.9, and 31.4. **(B)** Heatmap illustrating amplicon depth coverage across the same Ct values. The text on the y-axis lists the M-MLV RTase (wf-2) and the various polymerases (wf-3) evaluated in the study. The x-axis text highlight the amplicon IDs (1 to 99) from the SARS-CoV-2 genome targeted by the ARTIC primer scheme v4.1. Colour gradient: red (low values) indicates poor or no coverage (depth ≤ 20), representing amplicon dropouts.

In wf-3, where polymerases were compared, Platinum and EcoDry demonstrated the most consistent performance, closely matching the LoCost workflow with minimal dropout rates (Figures 2 and 3; *Extended data:* Table S4). Specifically, Platinum was the most comparable, achieving high genome coverage (97.3% to 99.6%) with only three amplicon dropouts. In contrast, under the conditions used, PrimeSTAR and Phusion were less reliable, with lower coverage (76.2% and 74.1%, respectively) at the highest Ct 31.4. We noted that the amplicons which failed to amplify efficiently varied between polymerases, with Platinum consistently exhibiting dropout of amplicon 90 and EcoDry primarily at amplicon 82 (Figure 2B). Amplicons such as 21, 22, 51, 60, and 66 frequently appeared as problematic areas across multiple conditions (including M-MLV RT with Q5 polymerase), particularly with high Ct samples. The results of wf-3 clearly demonstrated that Platinum and EcoDry polymerases can reliably substitute for Q5 polymerase in the LoCost workflow, maintaining high and stable genome coverage (96-99%) across all Ct values.

In wf-4, we replaced the commercial end-prep enzyme mix with generic HELP-EP mixes (A, B, and C), combining T4 PNK, T4 DNA Polymerase, Klenow, and Taq DNA Polymerase in a single reaction for amplicon end repair (Extended data: Table S1). HELP EP-A used the highest enzyme concentrations: 0.2 U/μl T4 PNK, 0.01 U/μl Klenow, 0.02 U/μl T4 Pol, and 0.04 U/μl Taq Pol. While HELP EP-B included 2-fold reductions in all enzymes except Klenow, which remained constant. HELP EP-C mirrored EP-A in enzyme concentration, but excluded Klenow to test whether T4 DNA Polymerase alone could support efficient end-preparation. Buffer components, 1X T4 DNA Ligase Reaction Buffer (1 mM ATP), 5% PEG-8000, and 0.5 mM dNTPs, were consistent across all HELP-EP mixes. As with wf1-3, all other steps of the sequencing workflow were maintained as per the LoCost protocol and the sequencing performance of each HELP-EP mix was compared using a dilution series of RNA extracted from the lab-grown SARS-CoV-2 isolate. HELP EP-C, excluding Klenow, was the most comparable to the LoCost protocol, achieving consistent genome coverage >95% across all Ct values (Figures 2 and 4; *Extended data:* Table S4). In contrast, HELP EP-A and HELP EP-B, while effective at lower Ct values, showed slightly reduced performance at higher Ct values (31.4), with genome coverage of 93.8%.

**Figure 4.**
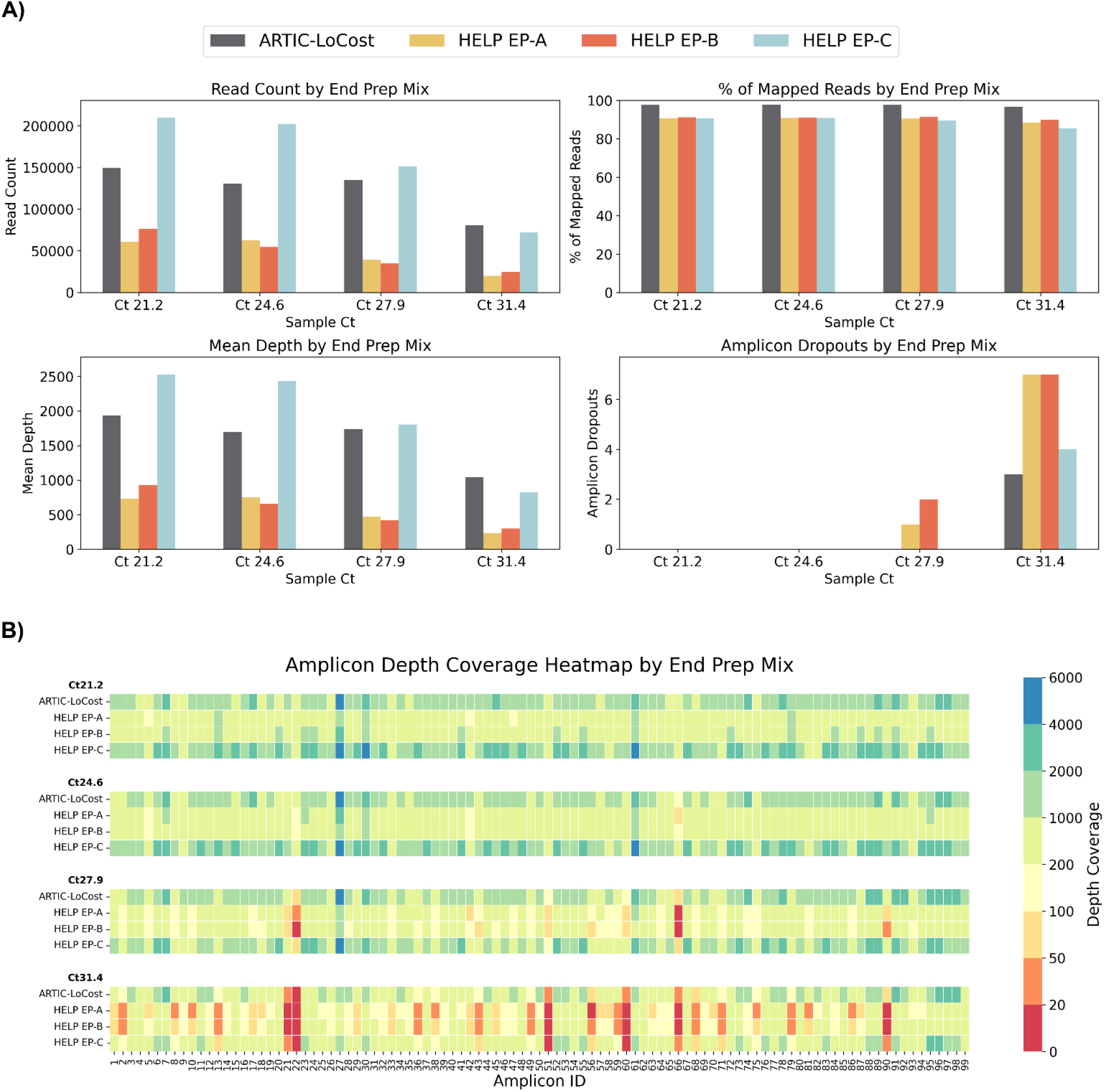
Stage 1: Comparison of HELP end-prep mixes to the LoCost protocol. **(A)** Bar plots comparing the performance of three HELP end-prep mixes (EP-A, EP-B, and EP-C) (wf-4) against the LoCost protocol (wf-1) across four key metrics: Read Count, % of Mapped Reads, Mean Depth, and Amplicon Dropouts. Evaluations were conducted using a SARS-CoV-2 isolate at Ct values of 21.2, 24.6, 27.9, and 31.4. **(B)** Heatmap illustrating amplicon depth coverage across the same Ct values. The y-axis lists the HELP EP mixes evaluated in the study. The x-axis text represents amplicon IDs (1 to 99) from the SARS-CoV-2 genome targeted by the ARTIC primer scheme v4.1. Colour gradient: red (low values) indicates poor or no coverage (depth ≤ 20), representing amplicon dropouts.

In wf-5, we evaluated five barcode ligation mixes (HELP BL, A–E), alongside a commercial DNA Ligation Kit, Mighty Mix (TAKARA) (Figures 2 and 5; *Extended data:* Table S4). Each HELP mix contained a consistent concentration of T4 DNA Ligase (1000 U/reaction) but differed in ligation enhancer supplements, including one or combinations of PEG-8000, 1,2-PrD, and HCC. HELP BL-D, which included both 10% PEG-8000 and 1 mM HCC, performed identically to the LoCost, achieving high genome coverage (>97%) across all Ct values. HELP BL-A (10% PEG-8000) and HELP BL-E (10% PEG-8000 with 2X Ligase buffer) also performed well, achieving genome coverage >95%. In contrast, HELP BL-B (15% PEG-8000) and HELP BL-C (10% PEG-8000 with 12% 1,2-PrD) had high genome coverage (>99%) at lower Ct values, but dropped to 93.8% at Ct 31.4. In contrast to the generic enzyme mixes, under the conditions used, Mighty Mix consistently showed the lowest performance across all metrics, with genome coverage dropping from 99.6% at Ct 24.6 to 97% at Ct 21.2 and 27.9, and further declining to 87.6% with 14 amplicon dropouts at Ct 31.4.

**Figure 5.**
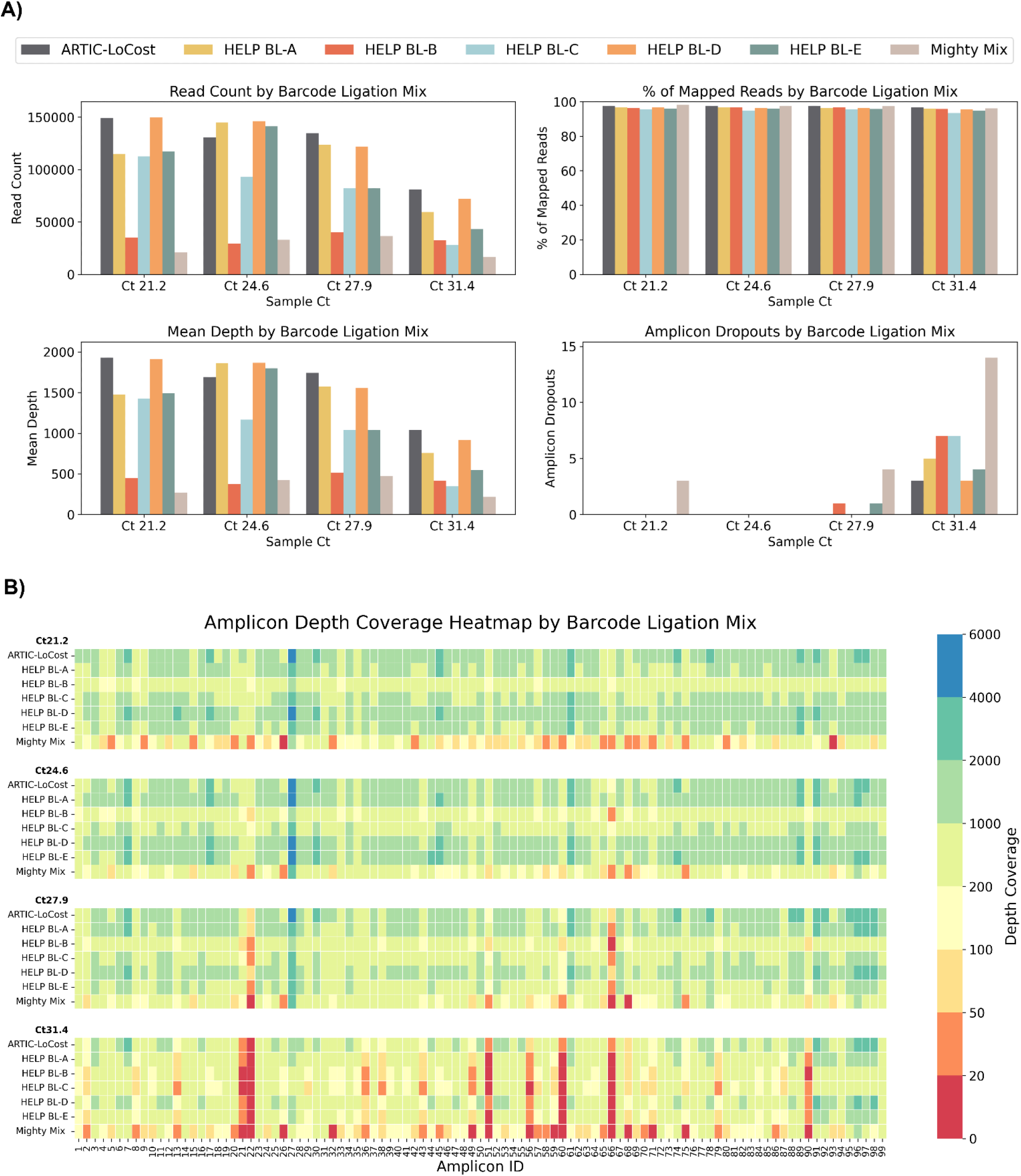
Stage 1: Comparison of HELP barcode ligation mixes to the LoCost protocol. **(A)** Bar plots comparing the performance of five HELP barcode ligation mixes (BL-A to BL-E) and a commercial Ligation Mighty Mix (TAKARA) (wf-5) against the LoCost protocol (wf-1) across four key metrics: Read Count, % of Mapped Reads, Mean Depth, and Amplicon Dropouts. Evaluations were conducted using SARS-CoV-2 isolate at Ct values of 21.2, 24.6, 27.9, and 31.4. **(B)** Heatmap illustrating amplicon depth coverage across the same Ct values. The y-axis lists the HELP BL mixes evaluated in the study. The x-axis represents amplicon IDs (1 to 99) from the SARS-CoV-2 genome targeted by the ARTIC primer scheme v4.1. Colour gradient: red (low values) indicates poor or no coverage (depth ≤ 20), representing amplicon dropouts.

In wf-6, we evaluated alternatives to the NEBNext Quick Ligation Module for adapter ligation in the ARTIC LoCost protocol by testing the HELP AL-A mix and Thermo Rapid DNA Ligation Kit (RLK, Thermo). The HELP AL-A mix was prepared with a final concentration of 2X T4 DNA Ligase Reaction Buffer, enhanced with 10% PEG-8000 and a high concentration of T4 DNA Ligase (4000 U/reaction). Both mixes showed performance comparable to the ARTIC LoCost workflow and proved to be reliable options for sequencing (Figures 2 and 6; *Extended data:* Table S4).

**Figure 6.**
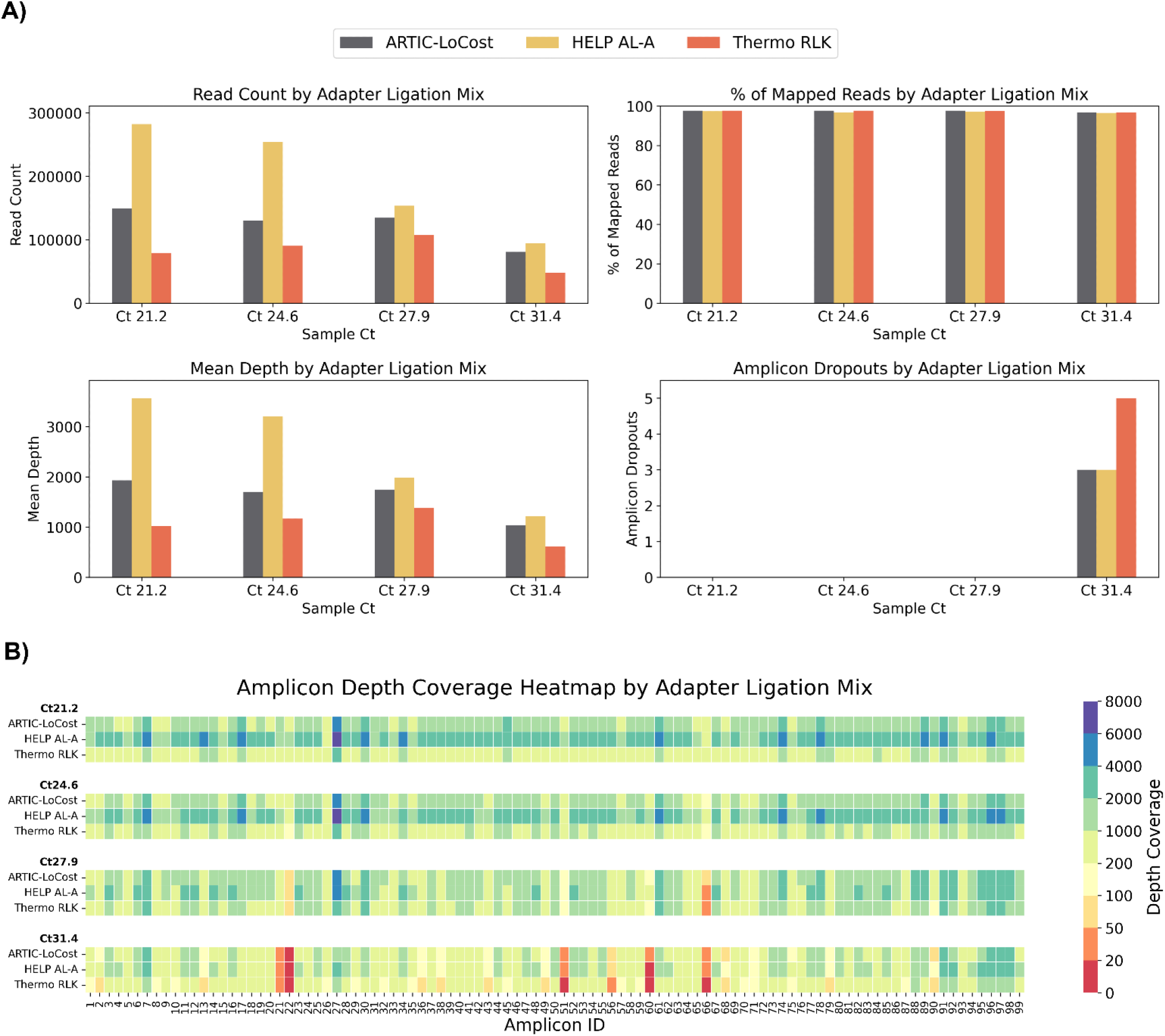
Stage 1: Comparison of HELP adapter ligation mixes to the LoCost protocol. **(A)** Bar plots comparing the performance of the HELP adapter ligation mix and the commercial Thermo Rapid DNA Ligation Kit (RLK) (wf-6) against the LoCost protocol (wf-1) across four key metrics: Read Count, % of Mapped Reads, Mean Depth, and Amplicon Dropouts. Evaluations were conducted using SARS-CoV-2 isolate at Ct values of 21.2, 24.6, 27.9, and 31.4. **(B)** Heatmap illustrating amplicon depth coverage across the same Ct values. The y-axis lists the mixes evaluated for adapter ligation (wf-6) in this study. The x-axis represents amplicon IDs (1 to 99) from the SARS-CoV-2 genome targeted by the ARTIC primer scheme v4.1. Colour gradient: red (low values) indicates poor or no coverage (depth ≤ 20), representing amplicon dropouts.

### Stage 2. Comparative Analysis of HELP Workflows Relative to the ARTIC LoCost Workflow

We next evaluated the performance of the generic replacements identified in Stage 1, within a single-library workflow, selecting only those that performed well in the initial screen. M-MLV served as a generic RT replacement, and for the PCR step, we selected Platinum and EcoDry polymerases due to their highest efficiency under the conditions used. For end prep, we selected HELP EP-A and EP-C mixes, which differed only by the inclusion of the Klenow enzyme in the EP-A. For barcode ligation, we selected HELP BL-A (10% PEG-8000) as the simplest recipe and BL-D (10% PEG-8000 and 1mM HCC) as the best performing mix. In the adaptor ligation step, we tested AL-A and AL-B which varied only by the amount of ligase per reaction, 4000U and 2000U, respectively. Stage 2 comprised two groups: Group I used Platinum polymerase and Group II using EcoDry polymerase, each with five workflows as illustrated in Figure 7. All combinations of the HELP workflows yielded results comparable to the ARTIC LoCost approach, although their performance varied across different sequencing quality control metrics, particularly as viral load in samples reduced (Figures 7 and 8; *Extended data:* Table S4).

**Figure 7.**
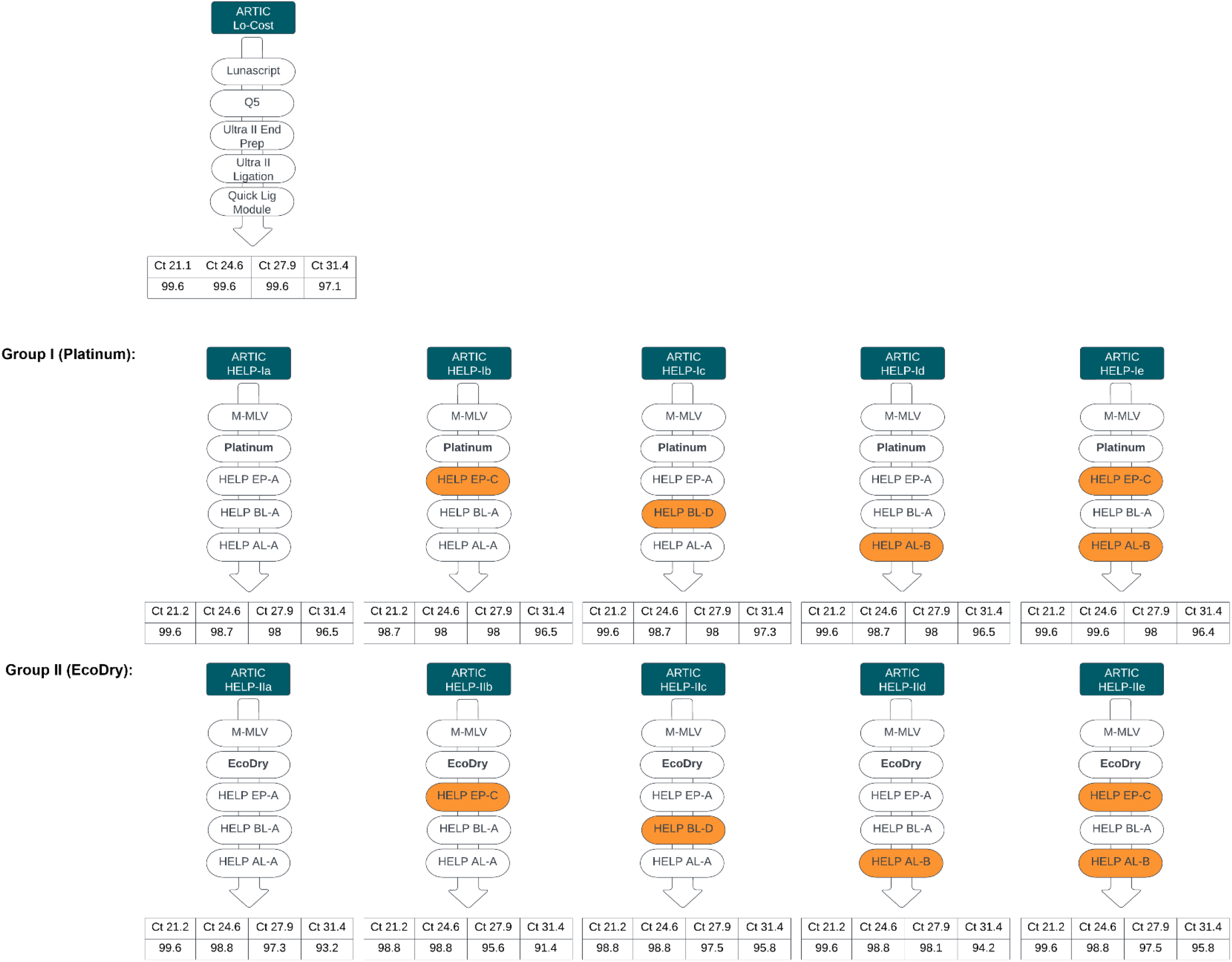
Stage 2: Summary of HELP workflow comparisons. Comparison of alternative HELP workflows (Groups I and II) with the LoCost protocol. Performance was evaluated across Ct values of 21.2, 24.6, 27.9, and 31.4 for SARS-CoV-2 isolate. **(Top)** Group I: HELP workflows Ia to Ie used M-MLV RT and Platinum polymerase for PCR, with variations in end-prep, barcode ligation, and adapter ligation mixes. **(Bottom)** Group II: HELP workflows IIa to IIe used M-MLV RT and EcoDry polymerase with the same variations.

**Figure 8.**
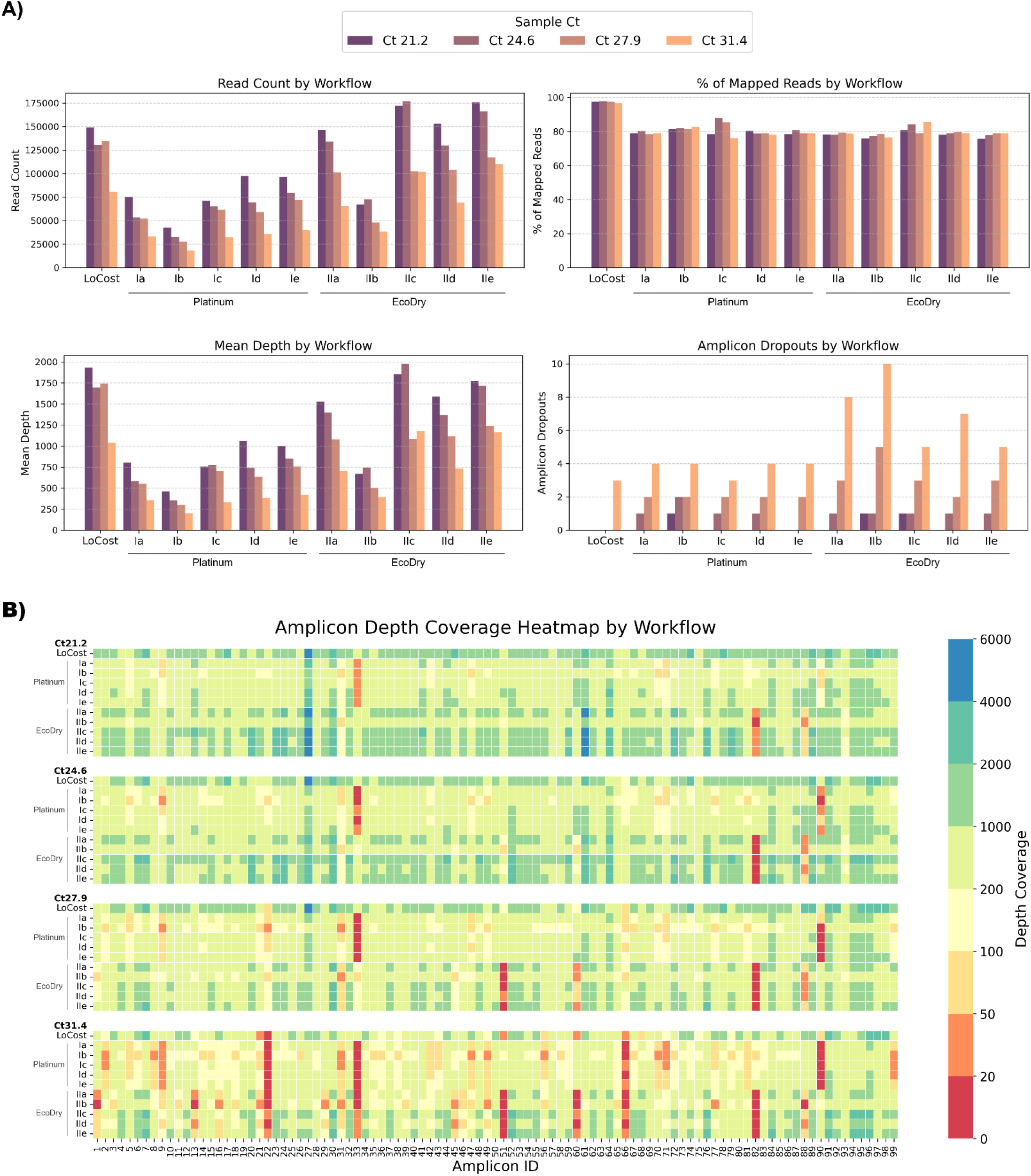
Stage 2: Performance of multiple HELP workflows compared to the LoCost protocol. **(A)** Bar plots comparing the performance of HELP workflows using two polymerase groups (Platinum and EcoDry) against the LoCost protocol across four key metrics: Read Count, % of Mapped Reads, Mean Depth, and Amplicon Dropouts, using SARS-CoV-2 isolate at Ct values of 21.2, 24.6, 27.9, and 31.4. Group I (Platinum): HELP-Ia to HELP-Ie workflows incorporating M-MLV RT and Platinum polymerase with variations in end prep, barcode ligation, and adapter ligation. Group II (EcoDry): HELP-IIa to HELP-IIe workflows using M-MLV RT and EcoDry polymerase with the same variations. **(B)** Heatmap illustrating amplicon depth coverage across various workflows. The x-axis represents amplicon IDs (1 to 99) from the SARS-CoV-2 genome targeted by the ARTIC primer scheme v4.1. Colour gradient: red (low values) indicates poor or no coverage (depth ≤ 20), representing amplicon dropouts.

Among the Platinum HELP workflows, HELP-Id and HELP-Ie showed the most consistent performance across all Ct values, making them the closest alternative to the LoCost. The remaining Platinum HELP workflows (Ia, Ib, and Ic) had similar percentages of mapped reads and genome coverage but exhibited more variability in mean depth. Specific dropouts were consistent at amplicon 33 and 90 across all Ct values for all Platinum workflows. Among the EcoDry HELP workflows, HELP-IId and IIe demonstrated relatively stable performance, making them good alternatives, though some dropouts were seen at higher Ct values. The remaining EcoDry HELP workflows (IIa, IIb, and IIc) showed more variability in mean depth and genome coverage, particularly at higher Ct values. Consistent dropouts were observed at amplicon 51 and 82 across all Ct values for all EcoDry workflows.

The results from the Stage 2 further confirmed the impact of polymerase choice and HELP mix combinations on sequencing yield. In general, HELP workflows using Platinum polymerase in all tested combinations consistently achieved genome coverage >95%, with minimal dropouts. In comparison, HELP workflows with EcoDry polymerase exhibited genome coverage ranging from >91%, depending on the HELP mix combination used.

### Stage 3. Validation of the HELP-workflow using clinical samples positive for SARS-CoV-2

To explore the utility of the best performing HELP workflows, HELP-Ie (Platinum) and HELP-IIe (EcoDry) workflows were compared to ARTIC LoCost with 19 SARS-CoV-2 (Delta variant) RNA clinical samples (Ct 20–33). Sample characteristics and comprehensive sequencing metrics are summarized in Extended data Table S5. For low Ct samples (≤24), LoCost and HELP-Ie outperformed HELP-IIe in genome coverage (98.1–99.6% vs. 95.6–98.2%) and fewer dropouts (Figure 9). At moderate Ct values (>24 to ≤28), HELP-Ie matched LoCost (95.5–98.9%), while HELP-IIe showed reduced performance (84.0–94.9%). At high Ct values (>28), HELP-Ie excelled (82.9–97.3%), LoCost showed moderate coverage (86.0–89.8%), and HELP-IIe performed worst (55.7–84.1%).

**Figure 9.**
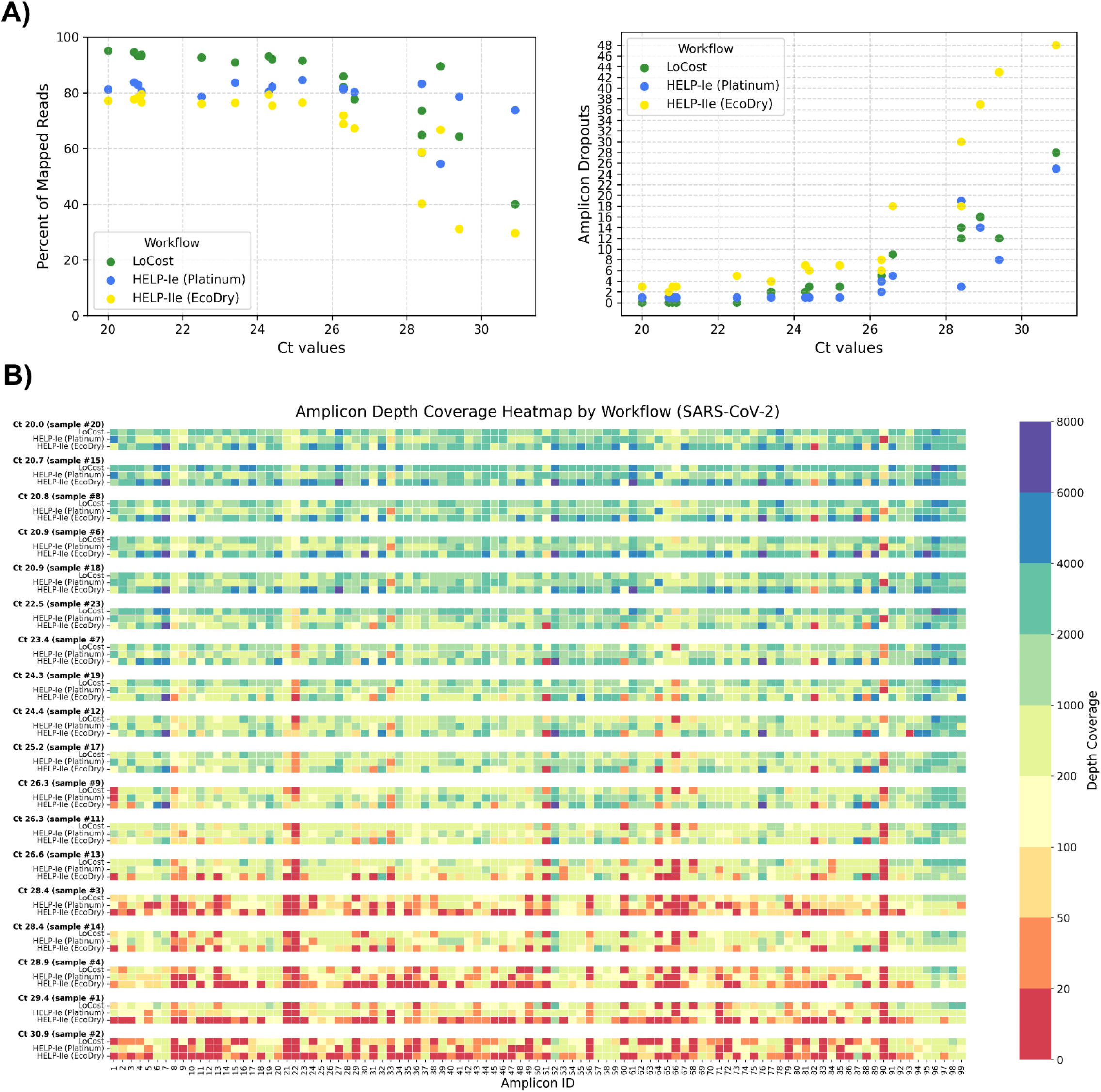
Stage 3: Performance on SARS-CoV-2 clinical samples. **(A)** Scatter plots showing the relationship between Ct values and (*left*) % of mapped reads and (*right*) amplicon dropouts across three workflows: LoCost (green), HELP-Ie (Platinum, blue), and HELP-IIe (EcoDry, yellow) using SARS-CoV-2 clinical samples. **(B)** Heatmap illustrating amplicon depth coverage for 18 SARS-CoV-2 clinical samples across workflows. Colour gradient: red (low values) indicates poor or no coverage (depth ≤ 20).

We also examined the amplicon dropout patterns using clinical samples (Figure 9B). Across all Ct values, specific amplicons (e.g., 21, 22, 66, 90) frequently showed dropouts across all workflows, particularly at higher Ct values. Additionally, each workflow exhibited a unique pattern of amplicon dropouts, suggesting that it may be possible to improve performance by rebalancing primer concentrations for the poorly performing amplicons (*30*). Amplicons 33 and 90 were determined to be unique to the Platinum polymerase, whereas for the LoCost workflow with Q5 polymerase, amplicons 22, 66, and 90 were the most sensitive sites. The EcoDry exhibited specific amplicon dropouts, such as 31, 51, and 82.

### Stage 3. Validation of the HELP-workflow using clinical samples positive for Norovirus GII

To confirm the potential applicability of the HELP workflow to viruses beyond SARS-CoV-2, we sequenced 12 Norovirus genogroup II-positive stool samples comprising eight different genotypes (Ct values 15.6–32.8) (*Extended data:* Table S6). Ten samples were unique, and two were included as technical replicates to complete a 12-barcode pool for optimal sequencing yield and barcode performance. We compared HELP-Ie (Platinum) and HELP-IIIe (Q5) workflows (substituting EcoDry with Q5) against the LoCost workflow. Q5, a benchmark polymerase for high-multiplex tiling PCR, was included to test its integration with the complete HELP workflow, spanning EP, BL, and AL steps. The results demonstrated that genome coverage and amplicon dropouts were predominantly influenced by genotype specificity rather than workflow or Ct values, highlighting areas for potential primer design improvement (Figure 10). Among the genotypes GII4P31, GII17P17, GII4P4 (recombinant), and GII4P16 exhibited no amplicon dropouts across all workflows, achieving genome coverage between 95.1% and 96.6% (excluding 5’ and 3’ ends as the primer scheme does not cover these regions). GII4P4 (Ct 24.9) and GII3P12 (Ct 15.6) showed moderate genome coverage (77.3–86.1%) with specific amplicon dropouts. GII6P7 (Ct 21.5) and GII7P7 (Ct 24.4) exhibited the lowest coverage (59.6–65.8% and 20.8–31.0%, respectively) with multiple consistent dropouts. These results demonstrated that the HELP workflow is robust and adaptable for sequencing Norovirus genogroup II, confirming its applicability beyond SARS-CoV-2. Coverage limitations observed for certain genotypes reflect primer scheme constraints rather than workflow performance, underscoring the need for further refinement of the norovirus-gii/800/v1.1.0 scheme.

**Figure 10.**
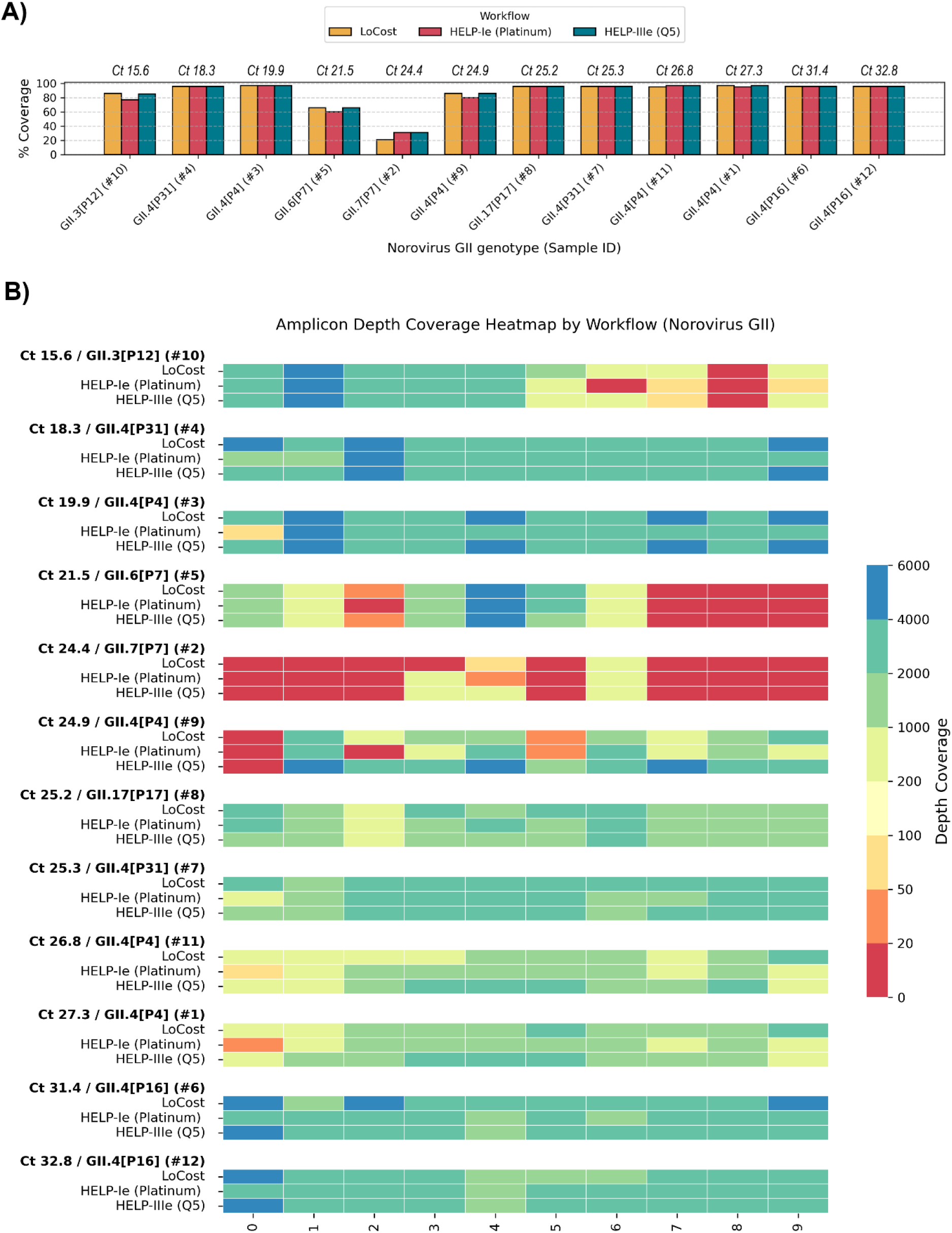
Stage 3: Performance on Norovirus GII clinical samples. **(A)** Genome coverage comparison across Norovirus GII genotypes and sequencing workflows. The sample Ct values are shown above the x-axis, with samples ordered from low to high Ct. Coverage is compared between the LoCost protocol and two HELP workflows using different polymerases: Platinum (HELP-Ie) and Q5 (HELP-IIIe). **(B)** Heatmap illustrating amplicon depth coverage for Norovirus GII sequencing. The workflows compared include the LoCost and two HELP workflows, utilising two polymerases: Platinum and Q5, with samples ordered from low to high Ct. The x-axis represents the amplicon IDs (0 to 9) targeted by the norovirus-gii/800/v1.0.0 scheme. Colour gradient: red (low values) indicates poor or no coverage (depth ≤ 20).

### Comparison between the cost of the HELP and LoCost workflows

We compared reagent costs for HELP and LoCost protocols across six countries (UK, India, Mali, Indonesia, the Philippines, Burkina Faso), including only the enzymatic steps of the workflow: RT, PCR, and library preparation (EP, BL, AL). Quotes were generated in each country, including additional taxes and delivery charges, to determine the cost of purchasing the reagents required for both protocols to the respective national or academic laboratories. As shown in Table 1, total costs for the LoCost workflow ranged from £30 in the UK to £57.70 in Indonesia. For the HELP workflow (using Platinum or EcoDry polymerase), total costs varied from £12.70 to £33.00. These costs exclude RNA extraction, SPRI clean-up, and ONT sequencing kit and flow cell. Despite this, notable price discrepancies were observed between countries. Across countries where complete data were available, the HELP workflow reduced reagent costs by approximately 48% to 60% compared to LoCost, for example, 58% in the UK and 60% in India.

**Table 1.** Cost comparison of reagent protocols across countries. Reagent costs per reaction are shown for the HELP and LoCost protocols, broken down into RT-PCR (using Platinum [A] or EcoDry [B]) and final library preparation steps (end prep, barcode ligation, adapter ligation). Total costs per country are included where both RT-PCR and final step data were available. The Philippines and Burkina Faso provided quotes only for library prep steps.

| Steps | Protocol | UK | India | Mali | Indonesia | Philippines | Burkina Faso |
| --- | --- | --- | --- | --- | --- | --- | --- |
| RT and PCR | A | £4.4 | £5.9 | £4.3 | £14.1 | n/a | n/a |
|  | B | £5.9 | £7.6 | £4.7 | £17.3 | n/a | n/a |
|  | LoCost | £5.0 | £8.0 | £5.3 | £9.8 | n/a | n/a |
| EP, BL, AL | HELP | £8.3 | £12.5 | £9.5 | £15.7 | £14.0 | £16.9 |
|  | LoCost | £25.0 | £38.2 | £27.6 | £47.8 | £38.5 | £52.1 |
| Total | HELP A | £12.7 | £18.4 | £13.8 | £29.7 | - | - |
|  | HELP B | £14.2 | £20.1 | £14.2 | £33.0 | - | - |
|  | LoCost | £30.0 | £46.2 | £32.8 | £57.7 | - | - |

We also compared total workflow costs (including ONT reagents, extraction, and clean-up) using UK pricing across library sizes of 24, 48, and 96 samples (*Extended data:* Table S7).

## Discussion

The ARTIC HELP protocol described here provides a viable alternative to the ARTIC LoCost and other amplicon-based native barcoding workflows (ONT), demonstrating comparable performance across a range of conditions. By validating alternative reagents and enzymes already available in many molecular biology labs, HELP addresses supply chain issues exposed during the COVID-19 pandemic and helps maintain genomic surveillance during future disruptions. While the ARTIC LoCost protocol has proven its versatility and effectiveness in advancing genomic surveillance (*9-14, 31-34*), the development of the ARTIC HELP workflow further enhances these capabilities.

Incorporating alternative enzymes and home-brew reagents into molecular diagnostics and sequencing workflows offers a practical solution for maintaining viral sequencing capabilities during reagent shortages and public health emergencies. Matute et al. (*35*) and Page et al. (*36*) demonstrated reliable home-brew RNA extraction and RT-LAMP diagnostic methods for SARS-CoV-2 detection without dependence on commercial kits. Similarly, Ou et al. showed that nanopore sequencing using home-brew components can match the performance of standard commercial mixes (*37*). In line with these findings, our study validated the effectiveness of HELP mixes with clinical SARS-CoV-2 and Norovirus GII samples, achieving high genome coverage. The integration of HELP alternatives into the LoCost protocol were implemented through practical single-step substitutions (stage 1) or combined workflows (stages 2 and 3). For a combined workflow most similar to the LoCost in terms of performance and consistency, HELP-e with Platinum polymerase is recommended, particularly for its balance across metrics and lower dropout rates. This workflow is available on protocol.io https://protocols.io/view/artic-help-protocol-for-amplicon-based-viral-genom-gzsibx6cf.

Focusing on single-step substitutions, we noted a number of key findings related to the performance of the HELP mixes. The HELP EP-C mix, which excludes the Klenow fragment, performed comparably to the ARTIC LoCost workflow, aligning with previous findings by Carøe et al. that simplified single-tube enzyme combinations can improve efficiency and accuracy (*37*). While not directly tested in our study, recent work (*38*) demonstrated that even T4 Polymerase may be dispensable in end-prep reactions. In that study, a homebrew solution containing only T4 PNK and Taq Polymerase effectively replaced commercial end-repair modules, likely due to the proofreading activity of high-fidelity polymerases, which generate blunt-ended PCR products. The inclusion of PEG in HELP EP mixes to enhance enzyme activity also reflects strategies reported by Neiman et al., who used PEG and T4 Ligase buffer in a single reaction to achieve efficient library preparation (*39*). For barcode ligation, HELP BL-A containing 10 % PEG showed the highest efficiency, consistent with reports that this concentration improves ligation, while higher levels such as 15 %, tested in BL-B, inhibited performance (*40*). HELP BL-C, which contained 1,2-PrD, demonstrated moderate efficiency but was less effective than HCC and PEG, which is in line with studies reporting 14-fold improvements with 1,2-PrD, and even greater gains, up to 50-and 100-fold, with HCC and PEG respectively (*40-43*). The highest performance was observed with HELP BL-D, which combined 10 % PEG and 1 mM HCC, suggesting a synergistic effect. Overall, the omission of Klenow in EP-C, the use of a simplified barcode ligation mix in BL-A, and a reduced ligase concentration in adaptor ligation mix AL-B were sufficient for effective performance in the combined HELP-e workflow. These findings reinforce the modularity and adaptability of the HELP mixes, allowing laboratories to customize workflows according to reagent availability and specific sequencing needs.

We found that the selection of the appropriate reverse transcriptase is particularly crucial at higher Ct values for optimal PCR efficiency. LunaScript, an engineered M-MLV RT variant, offers enhanced thermostability and reduced RNase H activity, leading to more efficient cDNA synthesis (*44*). In contrast, wild-type M-MLV RT, with its higher RNase H activity, can affect cDNA quality and yield (*45, 46*). While we showed that a wild-type M-MLV RT can serve as an alternative in the LoCost protocol when other options are unavailable, high-performance engineered reverse transcriptases, such as SuperScript IV, are recommended for better results, particularly in high Ct samples (>28).

Having multiple polymerase options provides additional flexibility to adapt to supply chain disruptions. High-multiplex PCR, such as the SARS-CoV-2 tiling scheme (*22*), uses around 50 primer pairs per pool to simultaneously amplify multiple regions of the genome, making polymerase selection critical for consistent results. This approach was originally optimised for Q5, a Pyrococcus-like (Pfu) B-family polymerase fused to the processivity-enhancing SSo7d domain, which provides high fidelity and strong template binding (*47*). In our study, all enzymes tested included proofreading activity, yet performance varied markedly depending on structural features and formulation. B-family Pfu-based polymerases with a SSo7d DNA-binding domain (e.g., Platinum, Phusion) performed well at low Ct values, but only Platinum sustained efficiency under low-input conditions, suggesting that even among similar polymerases, proprietary stabilisers and buffers influence template binding. Blended polymerases combining A-family (Taq-based) and B-family (Pfu-based) enzymes without an Sso7d DNA-binding domain (e.g., EcoDry, KAPA) showed variable performance. EcoDry consistently outperformed KAPA, potentially due to its lyophilised format and proprietary buffer composition. Notably, several polymerases that underperformed at higher Ct values still achieved strong coverage at Ct < 24, supporting their use as emergency alternatives for rapid outbreak-response sequencing when viral input is high and RNA quality is sufficient. These findings underscore that successful high-multiplex PCR depends not only on polymerase family or proofreading ability, but also on the presence of engineered DNA-binding domains and optimised buffer formulation, which together shape processivity, fidelity, and performance across varying template inputs.

We observed that amplicon dropout patterns in SARS-CoV-2 genome amplification varied by polymerase, even though most primers had well-matched melting (Tm) and annealing (Ta) temperature values, suggesting that Tm alone does not explain poor amplification. For example, amplicons 33 and 90 dropped out with Platinum, 51 and 82 with EcoDry, and 66 and 90 with Q5, all despite good predicted performance. It is likely that other factors, such as primer competition and differences in buffer composition, may have contributed to these dropouts. It is also notable that EcoDry recommends a higher annealing temperature (68°C), which may have affected its performance under the 63°C and 65°C conditions used in the ARTIC protocol with optimized primer concentrations and Tm. In contrast, Platinum and Q5 have broader Ta compatibility, making them more adaptable to these settings. Our findings align with previous studies of Lambisia et al., showing that rebalancing primer concentrations can help recover underperforming regions (*30*). While we did not test this directly, optimising primer pools per polymerase could improve consistency and should be considered.

We demonstrated that the HELP workflow is applicable to viruses beyond SARS-CoV-2 by sequencing Norovirus genogroup II. Our study also represents the first application of the norovirus-gii/800/v1.1.0 scheme (*23*), assessing its performance with both the LoCost and HELP workflows. Genome coverage varied across the eight Norovirus GII genotypes, likely due to primer mismatches driven by the high genetic diversity of GII noroviruses, rather than Ct values or workflows (*48, 49*). Still, we achieved genome coverage of >95% for four GII norovirus genotypes (GII3P12, GII4P4-recombinant, GII4P16, GII17P17), and >85% for two others (GII4P4, GII4P31). The lowest coverage was observed for GII6P7 (65.8%) and GII7P7 (31%), highlighting the need for improved primer design for these genotypes. There were also notable variations in mean depth for the same amplicon region across workflows (Figure 10B). Since primers were used in equal molar concentrations, our findings likely highlight the need for primer balancing for any given enzyme, to address these discrepancies and improve yield (*30*). To enhance amplification and coverage, targeted optimisations, such as adjusting concentrations of specific primers in pools, are recommended.

Our cost analysis highlights the well-established inequity in global access to molecular reagents (*50*). In some places, the same reagents cost nearly twice as much, with these cost increases arising not due to technical differences in the supplied products, but largely to customs fees, high import taxes, and shipping costs. This creates real challenges for labs in lower-resource settings, where budgets are limited and every extra cost makes it harder to carry out essential genomic work. The lack of transparency around the additional local added costs, how and when they are implemented, presents an additional barrier to the accurate budgeting of laboratory functions and grant proposals. These findings highlight the urgent need for more fair and transparent pricing, and support the value of locally sourced solutions like the ARTIC HELP protocol to make genomic research more accessible and affordable everywhere.

## Limitations of the study

Our study, while comprehensive, has limitations primarily due to the limited number of generic enzymes explored, the variability in polymerase performance and primer design. However we have provided a generic workflow template to identify other discrepancies in genome coverage and amplicon dropouts that challenge uniform amplification, particularly at high Ct values. Further optimization, including primer balancing and tailored reaction conditions for each polymerase, may be necessary to enhance the reliability and robustness of the sequencing workflow. Our results suggest that the norovirus-gii/800/v1.1.0 scheme requires improvements, particularly by designing additional primers that are more specific to GII6P7 and GII7P7 genotypes. Although the current norovirus scheme is still under development, our findings provide insights for further optimisation.

## Conclusion

We developed ARTIC HELP, a practical workflow using alternative enzymes and home-brew buffers that works as well as the standard ARTIC LoCost protocol for sequencing viral genomes from clinical samples. The ARTIC HELP workflow offers a practical solution to supply chain disruptions, supporting the continuity of critical sequencing activities and expanding genomic surveillance and diagnostic capacity during public health emergencies.

## Acknowledgements

This work was funded by the Wellcome Trust ARTIC Network Collaborative Award (206298/B/17/Z) and the Wellcome Trust Award (313694/Z/24/Z).

## Data availability

Raw sequencing data (FASTQ) and consensus genomes (FASTA) are available under ENA Project PRJEB89721. This project contains the following underlying datasets:

- stage1_sars_cov_2_consensus – SARS-CoV-2 consensus sequences from workflows 1–6. Includes a control panel of four SARS-CoV-2 RNA concentrations derived from a lab-grown isolate (SARS-CoV-2/human/Liverpool/REMRQ001/2020).
- stage1_sars_cov_2_fastq-raw – Raw sequencing data (FASTQ) from workflows 1–6 for the same control panel.
- stage2_sars_cov_2_consensus – SARS-CoV-2 consensus sequences from a control panel of four RNA concentrations derived from the lab-grown isolate (SARS-CoV-2/human/Liverpool/REMRQ001/2020).
- stage2_sars_cov_2_fastq-raw – Raw sequencing data (FASTQ) corresponding to the Stage 2 control panel.
- stage3_sars_cov_2_consensus – SARS-CoV-2 consensus sequences from a clinical panel of 19 samples processed using three workflows: LoCost, HELP-Ie (Platinum), and HELP-IIe (EcoDry).
- stage3_sars_cov_2_fastq-raw – Raw sequencing data (FASTQ) for the same SARS-CoV-2 clinical panel.
- stage3_norovirus_gii_consensus – Norovirus GII consensus sequences from a clinical panel of 12 samples processed using three workflows: LoCost, HELP-Ie (Platinum), and HELP-IIIe (Q5).
- stage3_norovirus_gii_fastq-raw – Raw sequencing data (FASTQ) for the same Norovirus GII clinical panel.

## Extended data

HELP workflows from the Stage 3 are available on protocol.io https://protocols.io/view/artic-help-protocol-for-amplicon-based-viral-genom-gzsibx6cf

Extended data supporting this study are available at: https://github.com/AnyaKovalenko/ARTIC-HELP

**This repository includes the following extended data files**:

*ARTIC-HELP_Extended-data.xlsx* file. This Excel file includes:

- Table S1. HELP Master Mix Recipes.
- Table S2. Reagent List used in the HELP study.
- Table S3. Characteristics of polymerases used in this study (as provided by the supplier).
- Table S4. Sequencing quality control metrics (stages 1 and 2: lab-grown isolate SARS-CoV-2/human/Liverpool/REMRQ001/2020).
- Table S5. Sequencing quality control metrics (stage 3: 19 clinical samples SARS-CoV-2).
- Table S6. Sequencing quality control metrics (stage 3: 10 clinical samples Norovirus Genogroup II).
- Table S7. Cost comparison per sample for libraries containing 24, 48, and 96 samples.

*ARTIC-HELP_Figure_S1.png* file :

Figure S1. Stage 1: Gel electrophoresis results for M-MLV (wf-2) and six polymerases (wf-3) compared to ARTIC LoCost (wf-1).

